# Decoding object categories from EEG during free viewing reveals early information evolution compared to passive viewing

**DOI:** 10.1101/2023.06.28.546397

**Authors:** Auerbach-Asch Carmel R., Vishne Gal, Wertheimer Oded, Deouell Leon Y.

## Abstract

Object processing is fundamental to visual perception, and understanding its neural substrates informs many cognitive and computational visual processing models. Thus far, most human studies have used passive viewing paradigms, during which self-driven behavior, such as eye movements, is constrained, and brain activity is evoked by abrupt stimuli onsets. This artificial dissociation of perception and action ignores the natural dynamics of visual processing. Thus, conclusions based on such passive viewing paradigms may not apply to active vision. Here, we study the human neural correlates of category representations during active visual processing by time-locking EEG to self-driven fixations during visual search for natural objects. We combine the deconvolution of overlapping responses to consecutive fixations with multivariate pattern analysis (MVPA) to decode object categories from responses to single fixation. We bridge the active and passive viewing literature by comparing the temporal dynamics of multivariate object representations during free visual search (active viewing) and rapid serial visual presentation (passive viewing), leveraging the high temporal resolution of EEG. We found that categorical information, at different levels of abstraction, can be decoded from single fixations during natural visual processing, and cross-condition decoding revealed that object representations are similar between active and passive viewing conditions. However, representational dynamics emerge significantly earlier in active compared to passive conditions, likely due to the availability of predictive information in free viewing. We highlight methodological considerations for combining MVPA with deconvolution methods.

**Significance Statement:** Understanding the neural correlates of visual perception is crucial for advancing cognitive and computational models of human vision. This study bridges the gap between passive- and active-vision literature while shedding light on the intricate relationship between perception and action in visual processing. Although eye movements are a fundamental behavior through which visual information is naturally sampled, most neuroimaging studies probe the brain by presenting stimuli abruptly at the center of the screen while participants refrain from moving their eyes. We investigated EEG correlates of visual processing during active visual search and demonstrated that object categories of naturally fixated objects can be decoded from the EEG. We provide novel findings regarding the dynamics of active, compared to passive, visual processing, while contributing to the advancement of EEG analysis methodology.

## Introduction

Object categorization is an essential component of visual perception (Rosch et al., 1976; Bracci and Op de Beeck, 2023). Early human studies have used fMRI and M\EEG to reveal functional anatomical modules (Malach et al., 1995; Downing et al., 2001; Kanwisher and Yovel, 2006), as well as critical time windows for categorical processing (VanRullen and Thorpe, 2001; Riesenhuber and Poggio, 2002; Rossion and Jacques, 2011; Ganis et al., 2012; Mudrik et al., 2014). More recently, multivariate pattern analysis (MVPA) has emerged as a data-driven method for identifying distributed object representations, without adhering to strong anatomical modularity (Haxby et al., 2001; Murphy et al., 2011; Cichy et al., 2014; Ray et al., 2020). Using MVPA and fMRI data, Kriegeskorte et al., 2008, found that the similarity of activation patterns between different objects reflects human-labeled object categories, with a hierarchy from the abstract (animate, inanimate) to concrete (snakes, tomatoes) categories. MEG (Carlson et al., 2013; Cichy et al., 2014) and EEG (Kaneshiro et al., 2015) single-trial MVPA decoding of object categories suggest that the brain uses transient and dynamically-evolving representations, to encode exemplar-specific information earlier than abstract categories (∼100 ms vs ∼240 ms).

Most neuroimaging studies use passive viewing paradigms in which behavior and brain activity are locked to abrupt changes in external stimuli, while observers refrain from self-driven eye movements. Such paradigms control the spatiotemporal attributes of visual information, yet natural vision is not a passive task. Instead, visual information is serially sampled by sequential eye movements. As eye movements are self-generated, the timing of visual input can be anticipated, and upcoming information can be partially predicted based on the gist of the scene, as well as pre-saccadic parafoveal information. The artificial dissociation of action from perception in the lab questions the validity of findings relying on passive viewing. For example, passive viewing studies arguably lead computational models of vision to focus on feedforward processing mechanisms (Marr, 1982; Serre et al., 2007; DiCarlo et al., 2012). To overcome these limitations, studies of active sensing (Schroeder et al., 2010; Zweifel and Hartmann, 2020), have promoted hypotheses that incorporate ongoing action-perception loops and predictive coding (Ahissar and Assa, 2016; Parr and Friston, 2017; Parr et al., 2022).

A growing body of research is adopting active viewing paradigms in human M\EEG studies thanks to advances in both eye tracking and analytic tools for deconvolving overlapping brain responses and eye-movement-induced artifacts (Dandekar et al., 2012; Smith and Kutas, 2015a, 2015b; Ehinger and Dimigen, 2019). These studies focus on brain activity locked to eye fixation onset (fixation-related potentials, FRPs), rather than on externally-evoked activity (event-related potentials, ERPs; Dimigen et al., 2011; Kamienkowski et al., 2012; Brouwer et al., 2013; Fischer et al., 2013; Frey et al., 2013; Körner et al., 2014; Devillez et al., 2015; Auerbach-Asch et al., 2020). Such natural paradigms reveal the extent to which pre-saccadic, extrafoveal information (Niefind and Dimigen, 2016; Buonocore et al., 2020; Huber-Huber et al., 2021), action-related corollary discharge (Cavanaugh et al., 2016; Sun and Goldberg, 2016), and temporal predictions can modulate foveal processing.

In the current study, we extend this burgeoning field by transitioning from the one-dimensional univariate analysis employed in active vision FRP research, to more powerful multivariate techniques, bridging the passive MVPA literature with active FRP literature (for limitations of univariate analysis, see Rousselet and Pernet, 2011). MVPA enables us to avoid pre-selection of spatial and temporal components of interest and to exploit single trial information rather than mean responses. We examine the dynamics of object representations during natural vision by applying MVPA decoding to simultaneous EEG and eye-tracking data in humans, comparing active viewing (free visual search; FVS) to passive viewing (rapid serial visual presentation; RSVP). To do so, we develop a procedure to combine the deconvolution of overlapping eye-related responses (Ehinger and Dimigen, 2019) with MVPA (Treder, 2020), highlighting methodological considerations when decoding co-registered EEG and eye-movement data.

## Materials and Methods

### Participants

Twenty-four healthy adults with no reported neurological illness participated in the experiment (14 females and 10 males; age range 18-33 years, 22.5±3.8 mean±std). The participants had normal or corrected to normal visual acuity by their report. Informed consent was obtained from all participants, and they received either payment (∼$12 per hour) or class credit for their participation. The institutional ethics committee of the Hebrew University of Jerusalem approved this study.

### Stimuli

For comparability with the existing literature, images were obtained from a shared dataset (Grootswagers et al., 2019a) of colorful natural objects, abiding by a well-studied hierarchical structure (Kiani et al., 2007; Carlson et al., 2013; Cichy et al., 2014; Kaneshiro et al., 2015; Grootswagers et al., 2017). Objects were divided into six categories: Human Faces (HF, N=102), Animal Faces (AF, N=68), Natural Objects (NO, N=162), Handmade Objects (HO, N=277), Animal Bodies (AB, N=96) and Human Bodies (HB, N=65; Figure 1.c). All images were luminance-equated, their histograms were matched using the Matlab SHINE toolbox (Dal Ben, 2019), and they were resized to 3°X3° visual degrees. Images were organized into 200 sets of 12 objects. For the visual search condition, each set was arranged in a three-by-four grid on a gray background (RGB [128,128,128]), separated by six visual degrees between object centers (Figure 1b). The array occupied 75% of the screen width and height, leaving a large margin from the array to the screen edges. We added a random jitter for the location of each image on the grid between −0.66 and 0.66 visual degrees, vertically and horizontally, and a random orientation jitter for each image between −45 and 45 degrees. Each array contained 0-4 objects from each of the six categories. Category locations within the arrays were balanced across arrays. We used two object types as targets of search, each used in 100 arrays (see Experimental Procedure): Butterflies (taken from the AB category) and flowers (taken from the NO category). Twenty-five percent (25%) of the arrays contained three targets; 40% contained two targets; 25% contained one target; and the remaining 10% contained no targets. Target locations on the screen were balanced as best as possible across the arrays. Target exemplars were presented only once throughout the experiment, while distractor exemplars did not repeat within a single array but could repeat across arrays. There were 390 non-target items per object category, except for the first ten subjects, where due to a technical error, there was an under-representation of Animal Bodies (270 repetitions) and an over-representation of Natural Objects (570 repetitions). This does not bias the decoding results since the classification was conducted on balanced groups (see MVPA decoding); however, the univariate analysis of the AB category is slightly noisier than the others. The same arrays, in a different order, were used for all subjects (see Experimental Procedure).

**Figure 1.**
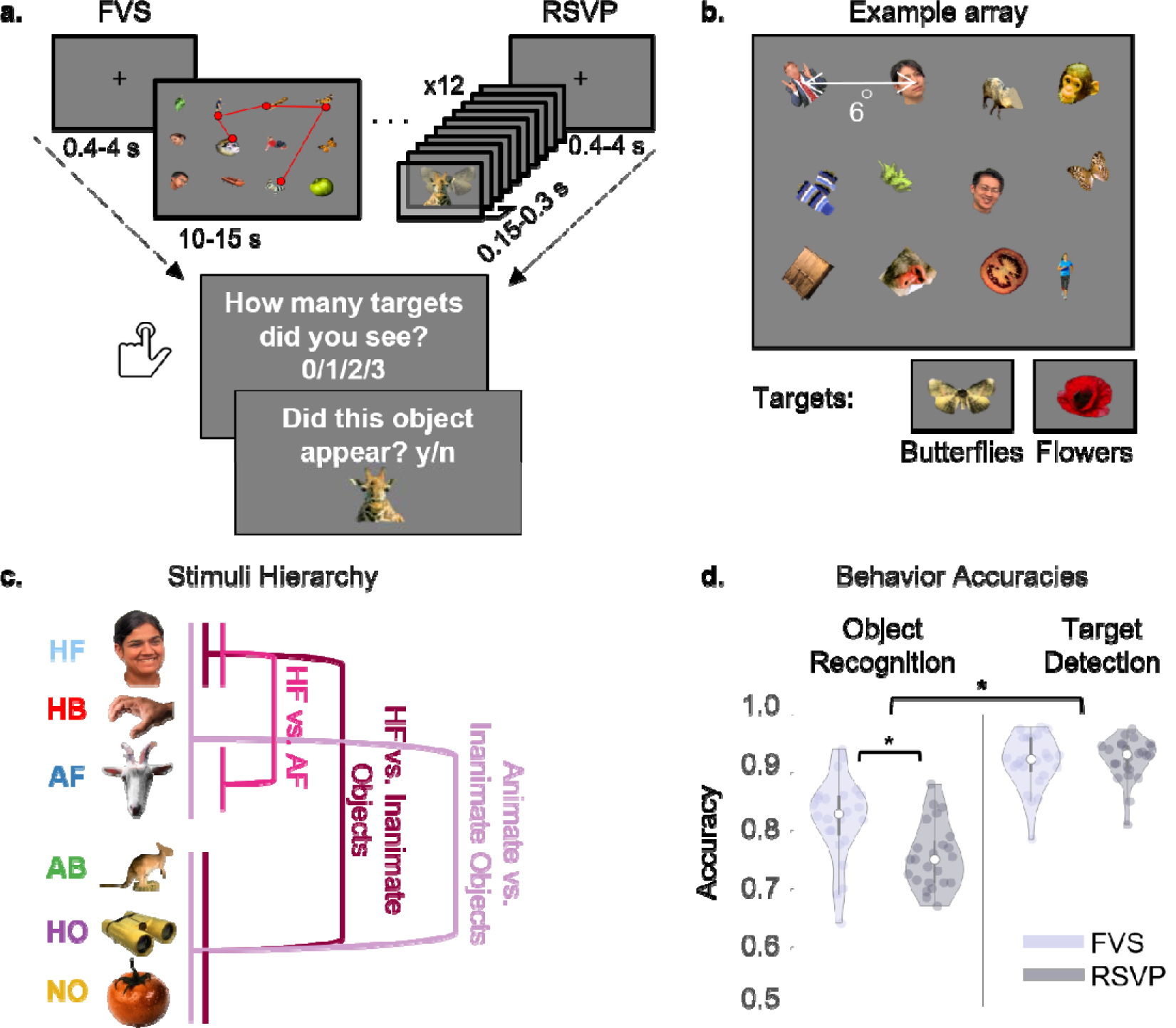
Experimental procedure and behavioral results. **a.** The two viewing conditions (presented in alternating order): Free Visual Search (FVS) - object sets were presented simultaneously on the screen in a search array. Rapid Serial Visual Presentation (RSVP) - objects were presented sequentially at the center of the screen in sets of 12. In both viewing conditions, after each set of objects, participants reported the number of targets detected and then performed an object recognition task. **b.** Top: example of a 12 objects array used in the FVS condition. Note that in the experiment, the array occupied 75% of the screen width and height; the borders were cropped in the figure for better visualization of the objects. Bottom: example of flower targets and butterfly targets (used in separate sessions). **c.** Exemplars from the six stimuli categories - Human Faces (HF), Human Bodies (HB), Animal Faces (AF), Animal Bodies (AB), Handmade Objects (HO), and Natural Objects (NO). The analysis was performed on three main categorical comparisons: HF vs. AF; HF vs. Inanimate Objects (IO); and Animate (A) vs. IO. Images obtained from an open dataset https://osf.io/a7knv/ **d.** Behavioral accuracies for the object recognition (left) and target detection (right) tasks in the FVS and RSVP conditions. The filled circles represent single subjects and the white circle represents the median across subjects. The violin contour represents the kernel density estimation of the data distribution; the thick vertical black line depicts the interquartile range of the data; and the thin vertical line depicts the full data distribution excluding outliers that exceed the 1.5*interquartile range. The asterisks mark significant effects.

### Experimental Procedure

Participants sat in a sound-attenuated room, 60 cm from a 50 X 30 cm screen (BENQ XL2411P) with a 144 Hz refresh rate and resolution of 1920 X 1080 pixels. The experiment included two conditions: FVS and RSVP (Figure 1a), each including 200 sets of 12 images. All sets began with a fixation cross at the center of the screen presented for 400-4000 ms (chosen randomly from a uniform distribution), upon which eye-tracking was validated (see EEG and Eye-tracking Coregistration). In the FVS condition, the image set was then presented as an array of 3X4 objects, and participants freely viewed the array in a self-paced manner for 15 seconds while their gaze position was tracked. In the RSVP condition, the gaze was restricted to the center of the screen, and the set of twelve objects was presented serially, centered on the fixation point, with no inter-stimulus interval. Stimulus Onset Asynchrony (SOA) was randomly and uniformly distributed between 100-300 ms (with 1 ms steps), with a mean of 200 ms, chosen to match the expected mode of fixation duration distributions (Findlay and Gilchrist, 2003). Each array in the FVS and each series in the RSVP included 0-3 flower or butterfly targets with the proportions described in the <u>Stimuli</u> section. In both viewing conditions, the set presentation ended with two, non-speeded behavioral tasks: target detection and exemplar recognition (always in this order).

In the target detection task, participants were required to report the number of targets presented in the preceding set. In the exemplar recognition task, participants were presented with a non-target object and were required to decide if that object was presented in the preceding set. In 50% of the sets, the object was taken from the preceding set, and in the remaining 50% the object was not present in the preceding set (but could have been shown in other sets). The recognition task was included to encourage participants to pay attention and fixate on all the objects rather than only on the targets.

The experiment was divided into two consecutive sessions separated by a ∼5 min break: one with butterflies as targets and one with flowers as targets. The order of the sessions was counterbalanced between subjects. Each session was split into 20 blocks, half RSVP and half FVS, alternating between viewing conditions. The condition of the first block was counterbalanced across subjects (i.e. ABABAB… or BABABA…). Each block consisted of 20 object sets, and blocks were separated by an optional break. At the start of each session, participants performed one training set for each viewing condition to familiarize them with the structure of the two conditions and with the session targets.

### EEG and Eye-Tracking Coregistration

EEG was acquired using an ActiveTwo system (Biosemi, The Netherlands) with 64 active electrodes mounted on an elastic cap according to the extended 10–20 system. The EEG sampling rate was 1024 Hz and an online low-pass filter with a cutoff of 1/5 of the sampling rate was applied to prevent aliasing. In addition, eight active, flat electrodes were placed: two on the mastoid processes, two horizontal electrooculogram (EOG) electrodes next to the outer canthi of the left and right eyes, two vertical EOG electrodes below and above the right eye, a single EOG channel under the left eye, and a channel on the tip of the nose. We referenced all electrodes during recording to a common-mode signal (CMS) electrode between POz and PO3. We recorded binocular eye movements using a desktop-mounted Eyelink 1000/2K infrared video-oculography system (SR Research Ltd., Ontario, Canada) at a sampling rate of 1000 Hz. A 12-point calibration procedure, followed by a validation stage (with acceptance criteria of worst point error < 1.5 visual degrees and an average error < 1.0 visual degrees), was applied before each block. We performed a gaze-drift check before every object-set presentation by ascertaining the registration of the eyes within a 3°x3° visual degrees square centered on the center fixation point for at least 400 ms. If this requirement was not reached within four seconds, the calibration procedure was repeated until this requirement was met.

EEG and eye-movement synchronization was performed offline using triggers sent via a parallel port from the stimulation computer, which were split and recorded simultaneously by the EEG recording system and the eye tracker. In addition to the digital eye-tracker data, we also sampled analog gaze position and pupil size from the eye tracker in the Biosemi system, simultaneously with the sampling of the EEG. We used this data to ensure accurate synchronization of EEG and eye movement data, but performed eye-movement analysis on the digital data, which has a better signal-to-noise ratio (SNR).

### Eye-Movement Preprocessing

We recorded binocular eye movements for 21 participants and monocular eye movements for three participants due to difficulty in tracking one of the eyes. We defined blinks as periods of simultaneous data loss in both eyes (or one eye in the three participants recorded monocularly). We defined saccades and fixations using an algorithm adapted from the EEGLab toolbox, which defines saccades as outliers in the two-dimensional eye velocity space (Engbert and Mergenthaler, 2006). We excluded detected saccades if they lasted for less than 10 ms, their amplitude exceeded the diagonal screen size (>30°), or their velocity was larger than 1000 ms/deg. Fixations were excluded if they lasted less than 30 ms or more than 1500 ms. On average,LJ7% (SEM = 1.2%) of fixations were removed per participant due to these exclusion criteria. The movement of the eyelid at the beginning and end of a blink may sometimes be classified by the above algorithm as a sharp vertical gaze shift. Therefore, we expanded blink windows to exclude such effects, opting for missing saccades over including spurious ones.

We defined Fixations of Interest (FOIs) as fixations landing within a radius of 1.5 visual degrees around the center of a single object in the search array (the region of interest; ROI). A valid First Fixation (FF) was defined as the first time, within each array, that the eyes fixated on an ROI of a certain object. Return Fixations (RFs) were defined as fixations landing on a previously fixated object immediately following a fixation on another object. “Ensuing Fixations” (EFs) were defined as fixations following either an FF or RF while dwelling within an object’s ROI. Any fixations landing between objects or on the periphery of the array were defined as General Fixations (GFs) and included in the analysis only to account for overlapping activity (see <u>Temporal Signal Deconvolution using GLM</u>).

### EEG Preprocessing

For data processing, we used custom Matlab code (Gerber, 2019), with some functionality adopted from the Fieldtrip toolbox (Oostenveld et al., 2011), EEGLab toolbox (Delorme and Makeig, 2004) and the Unfold toolbox (Ehinger and Dimigen, 2019). Electrodes that included excessive noise across the entire experiment duration (based on visual inspection) were excluded (ten participants lost one noisy channel, and two participants lost four channels due to channel bridging to the reference channel). Next, data were re-referenced to an average of all remaining electrodes. The recording was stopped between blocks. To concatenate the data to one continuous recording without amplitude step artifacts that would be later smeared by filtering, we applied linear detrending within each block by subtracting a linear vector connecting the means of the block’s initial and final ten samples. We next applied a third-degree, zero-phase-shift Butterworth high-pass filter on the data with a cutoff of 0.1 Hz (except for two participants with large low-frequency drifts due to sweating, for which we used a cutoff of 1.3 Hz). We removed 50 Hz line noise with a custom notch filter designed to suppress continuous 50 Hz oscillations, typical of power-line noise, and harmonics thereof, while having less effect on more transient components (Keren et al., 2010).

We applied independent component analysis (ICA) to attenuate noise driven by eye movements and muscle activity. For the detection of eye-movement-related ICA components, training data was generated by first removing large artifacts exceeding a threshold of 400 µV, followed by filtering between 2-100 Hz (using a 4th-degree non-causal Butterworth filter), and then concatenating segments of −200 to +1000 ms around stimuli onsets and segments of −30 to +30 ms around saccade onsets. These parameters were found to be optimal for training ICA for removing ocular artifacts (Keren et al., 2010; Dimigen, 2020). We manually identified ICA components as reflecting muscle, blink, or eye-movement artifacts, based on their temporal profile, scalp topography, and power spectrum averaged around blink, saccade, or stimulus onsets. The ratio of component variance during saccades and fixations was used as an additional measure for eye-movement-related artifacts (Plöchl et al., 2012; Dimigen, 2020). Selected components were removed from the original preprocessed data (to be distinguished from the ICA training data) and remixed into channel data (mean±SEM number of components removed per subject: 12±0.87). Next, time points in which activity exceeded a threshold of ±100 µV were marked as artifacts (with an additional margin of 40 ms before and after), and rare additional artifacts were marked following a visual inspection. All segments overlapping artifacts were excluded from the analysis. Finally, electrodes excluded before preprocessing were recreated by mean interpolation of the neighboring electrodes, and the data were downsampled to 256 Hz for further analysis.

### Combined Temporal Signal Decomposition and MVPA Decoding

Deconvolving temporally-overlapping EEG responses (such as in RSVP or free viewing) is commonly done using multiple regression (general linear model, GLM), which also enables accounting for continuous covariates such as saccade direction and amplitudes (Dandekar et al., 2012; Smith and Kutas, 2015a, 2015b; Nikolaev et al., 2016; Ehinger and Dimigen, 2019). This results in time-resolved estimates (beta coefficients) of the contribution of each modeled event separately, referred to as regression Fixation-Related Potentials (rFRPs) or regression Event-Related Potentials (rERPs), replacing traditional trial-averaged FRPs or ERPs. However, this approach only generates mean estimated responses. To obtain single-trial deconvolved data necessary for MVPA decoding, we reconstructed the single trials using estimates of the overlapping responses, as explained in the next section.

### Temporal Signal Deconvolution using a General Linear Model

We used GLM-based deconvolution (Smith and Kutas, 2015b; Ehinger and Dimigen, 2019) to obtain FRPs. A separate GLM was designed for the FVS and RSVP viewing conditions, including predictors corresponding to the relevant events; each predictor was modeled from 200 ms prior to the event to 1500 ms after the event. In the FVS condition, we defined separate binary predictors for three types of fixations (FF, RF, and EF, see <u>Eye Movement Preprocessing</u>) on each of the six categories of objects, resulting in 18 different fixation predictors. We added one binary predictor to account for any fixation that did not land within an object ROI (GF). In addition, we added a binary predictor to model the effect of array onset and two binary predictors to estimate the target-specific response to butterflies and flowers separately. Finally, each fixation predictor was accompanied by its incoming saccade-direction continuous predictor, split to its cosine and sine values, and a saccade-amplitude continuous predictor spread over a five splines basis set (Ehinger and Dimigen, 2019). The Wilkinson notation (Wilkinson and Rogers, 1973) of our GLM is:

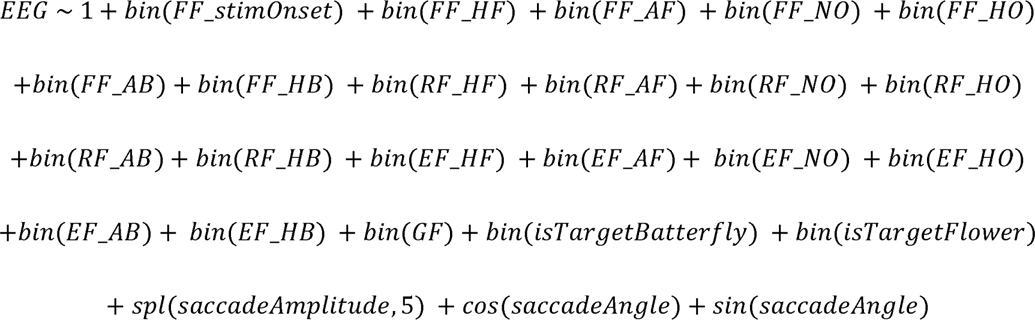

Where bin^1^ stands for a binary predictor, spl stands for spline basis, and cos\sin refer to the cosine and sine basis.

The GLM for the RSVP condition included one predictor set for the stimuli onsets of each of the six object categories listed above, one predictor for each of the two target categories, and a predictor for general fixations to account for eye movements occurring during RSVP:

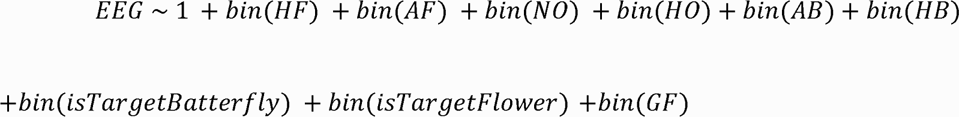

The design matrix was expanded to model each time point around each event separately during a window of −200 ms to 1500 ms around event onset and solved using the Matlab lsmr function (Fong and Saunders, 2011). For a detailed explanation of this step, the reader is referred to (Smith and Kutas, 2015a).

We used a data reconstruction approach to obtain single-trial deconvolved data (Figure 2a-d). For each event of interest, we used the mean rERP/FRP estimates of all proximal events to reconstruct the neural responses overlapping the event of interest. We then subtracted this reconstructed time course from the raw data before segmenting the isolated event of interest (Figure 2d). We refer to the single trials after overlap deconvolution as “GLM reconstructed single trials”. The effect of temporal signal deconvolution on both the single trials and the grand average responses, compared to classic averaging, is presented in Figure 2e.

**Figure 2.**
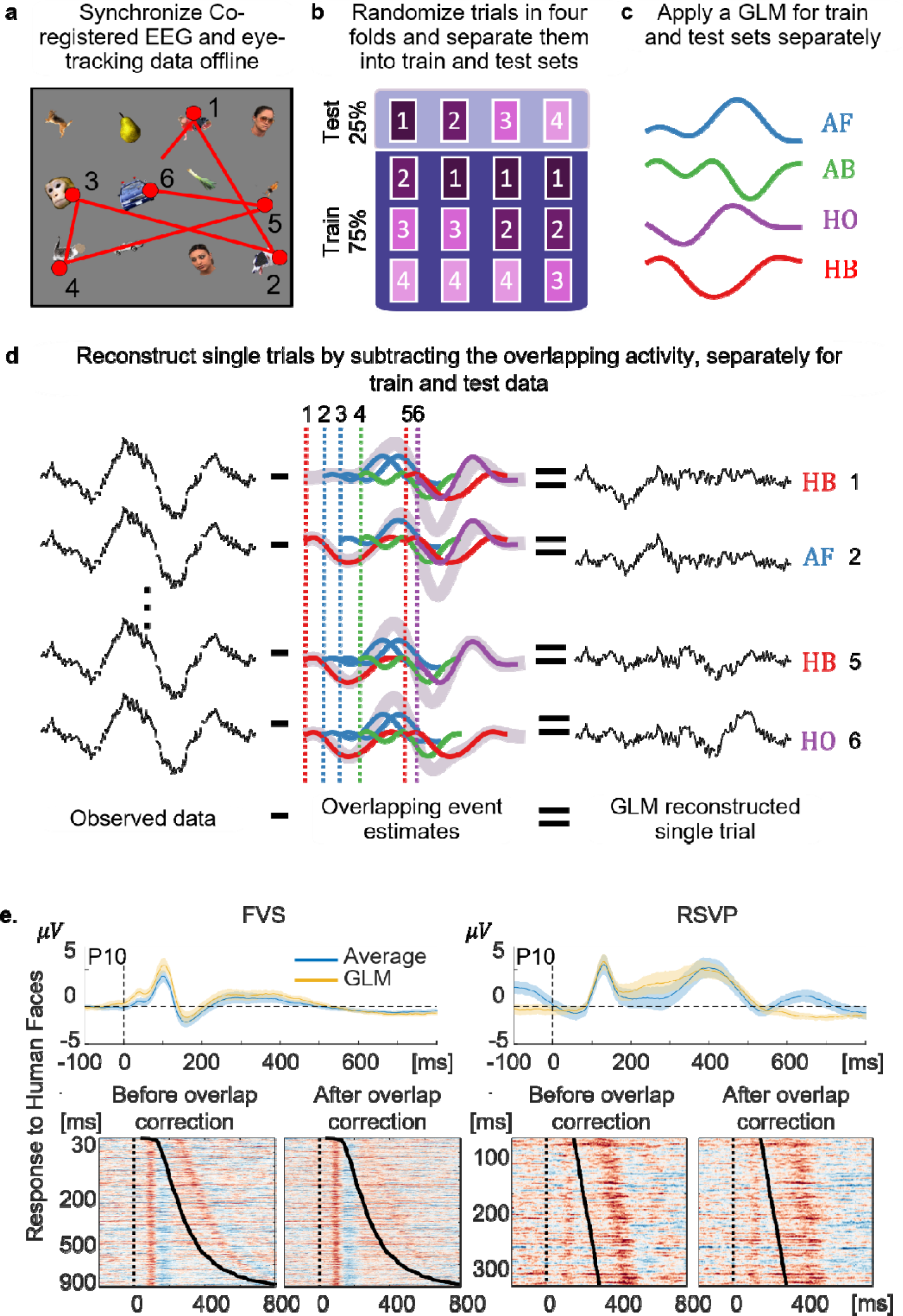
MVPA decoding after signal deconvolution. **a**. Simulated example of eye movements that define the sequential events of interest in the GLM (EOIs). For the illustration, we only show an example of first fixations (FF) on objects. **b**. Four-fold cross-validation: Trials were first randomized into four folds. Each fold was used once as a test set once (top), with the corresponding train set including data from all other folds (75-25% split; bottom). **c**. We obtained estimated mean responses after signal deconvolution for fixations on different categories, by applying a GLM to each train and test set separately. **d**. For each EOI (1-6 in the example illustration), overlapping responses from all events except of that EOI were reconstructed (the sum of all overlapping events is depicted by the thick gray line) and subtracted from the data, resulting in the estimated single trial response after overlap correction. **e.** Effect of signal decomposition on the data. Top: Average over subjects of the FRPs (left) and ERPs (right) obtained by classic trial averaging (blue) compared to those obtained by GLM reconstructed trials (yellow) in response to Human Faces in electrode P10. The shaded margins represent 95% confidence intervals across subjects. Bottom: Trials stacked and sorted by fixation duration in the FVS condition (left pair of graphs) and by stimulus duration in the RSVP condition (right pair of graphs) for one representative subject in response to Human Faces. For each condition, we compare trials before (left) and after (right) applying the GLM deconvolution of overlapping responses. AF - Animal Face, HF - Human Face, AB - Animal Body, HB - Human Body, HO - Handmade Objects.

During decoding, classifier weights are obtained using a training set, which must be distinct and independent from the test set used to quantify the classifier’s performance. In the present case, if rERPs\rFRPs are estimated using the entire dataset (as is typically done in univariate analyses), and then single trials of the train and test sets are reconstructed using the estimates based on the entire dataset, overfitting may occur (‘double-dipping’; Kriegeskorte et al., 2009). The GLM and reconstruction were therefore applied to training sets and testing sets of each fold separately.

### MVPA Decoding

To quantify the representation of category-related information locked to either stimulus onset (RSVP) or single fixations (FVS), we trained a set of linear classifiers to decode object categories from reconstructed single presentations or fixations on a time-point by time-point basis. To avoid overfitting, we applied a 4-fold cross-validation procedure; for each fold, 75% of the data was used as a training set, and the remaining 25% as the test set (Figure 2b). Importantly, as noted above, we obtained single-trial estimates by using the GLM approach separately for the train and test sets. The entire 4-fold procedure was repeated twice with different fold assignments. Across subjects, the number of fixations per category included in each fold in the FVS was FF: 353±36, RF: 437±50, and EF: 93±9 (Mean±SEM across participants). Only FF and RF types (combined) were used for category decoding. In each classification between two categories within a fold, the category containing more trials was undersampled to balance the number of trials between the two categories. Single trials were baseline corrected to the mean activity between −200 to −50 ms before stimulus or fixation onset.

For classification, we applied Linear Discriminant Analysis (LDA) using custom code based on the MVPA-Light toolbox (Treder, 2020). LDA attempts to find a linear weighting (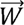) of the scalp electrodes for which separation between the categories is maximal. This is achieved by simultaneously maximizing the distance between categorical mean responses (“signal”) while minimizing response variability (“noise”), using the following formula (where 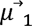 and 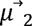 are the mean responses to each category and Σ: is the covariance matrix obtained by averaging the covariance matrices of each category separately):

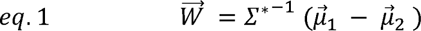

To reduce the influence of random noise fluctuations on the weight estimate, we used a shrinkage estimator for the covariance matrix (Σ:*; Blankertz et al., 2011) by combining the empirical covariance (Σ) and the identity matrix (*I*), weighted according to a regularization parameter (*λ*):

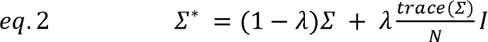

Multiplication of the identity matrix by the mean value on the covariance diagonal (N is the number of features of the classifier, equal to the number of electrodes in this case) ensures that the trace of the covariance is preserved, which helps to mitigate the bias introduced by this step. The regularization parameter *λ* was estimated using the Ledoit-Wolf formula (Ledoit and Wolf, 2004); implemented in MVPA-Light.

Once the linear weights (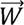) were obtained, the threshold for separating between the two categories was computed as:

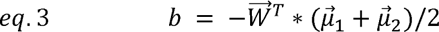

The predicted category for each test-set trial at each time point is determined by multiplying the vector of electrode activations (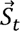) with the feature weights (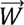) and adding the threshold term (*b*; eq. 3):

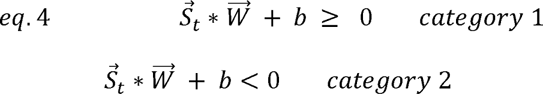

The accuracy of the classifier for each time point was defined as the percentage of correct classification it made on the test set for that time point. This was averaged across four folds and two repetitions of the cross-validation procedure.

We obtained decoding accuracies for each subcategory (Figure 4c) in two ways: 1. By classifying each category against all the other five categories (one vs. all, where the ‘all’ includes the same number of trials as the ‘one’, equally divided between the remaining five categories). 2. By classifying every pair of the six subcategories with each other (one vs. one), and for each category, averaging the results for pairs including that category (e.g HF vs. AF, HF vs. AB, etc.). The two methods produced similar results and we report the former (one vs. all) for the analysis.

For interpretability of the classifier weights in terms of activation patterns (or the extent to which each feature \ electrode contributed to the classification decision; Figure 4), we corrected the weights using the transformation suggested by (Haufe et al., 2014):

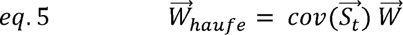

To test for representation stability, we used the temporal generalization method (King and Dehaene, 2014; King et al., 2014). If category representation is similar in time *t*_1_ and *t*_2_, the weights obtained from a classifier at time *t*_1_ will be able to classify data at time *t*_2_. Therefore, classifiers were used to classify all the time points in the test dataset (not only the time point used for the training) resulting in Temporal Generalization Matrices (TGMs; Figure 5c). In addition to decoding category representations within each viewing condition separately, we further performed cross-condition decoding. This is achieved by creating training sets from data obtained during active viewing (FVS) and testing classifier performance on a test set obtained during passive viewing (RSVP), and vice versa.

### Statistics

#### Behavioral Analyses

In the target detection task, participants were asked to indicate the number of targets presented in the previous set of objects (0-3). In the object recognition task, participants were required to indicate whether a specific exemplar was presented in the set or not (“yes”\”no”). We calculated response accuracies for each subject and task as the percentage of sets for which a correct response was provided. We ran a two-way, repeated-measures analysis of variance (ANOVA) to analyze the effect of viewing condition (passive or active), and task (target detection or object recognition) on the participants’ performance accuracy, collapsing over the two target categories (Figure 1d).

#### Eye-Movement Analysis

We tested whether there was a difference in eye movement parameters depending on the object category on which the eyes landed during the FVS condition. Eye movement parameters included: (1) amplitude of the saccade leading to the fixation; (2) direction of the saccade leading to the fixation; (3) fixation duration. For each of these parameters, we ran a one-way, repeated-measures ANOVA to test for differences in these measures depending on the object category (six levels; Figure 3).

**Figure 3.**
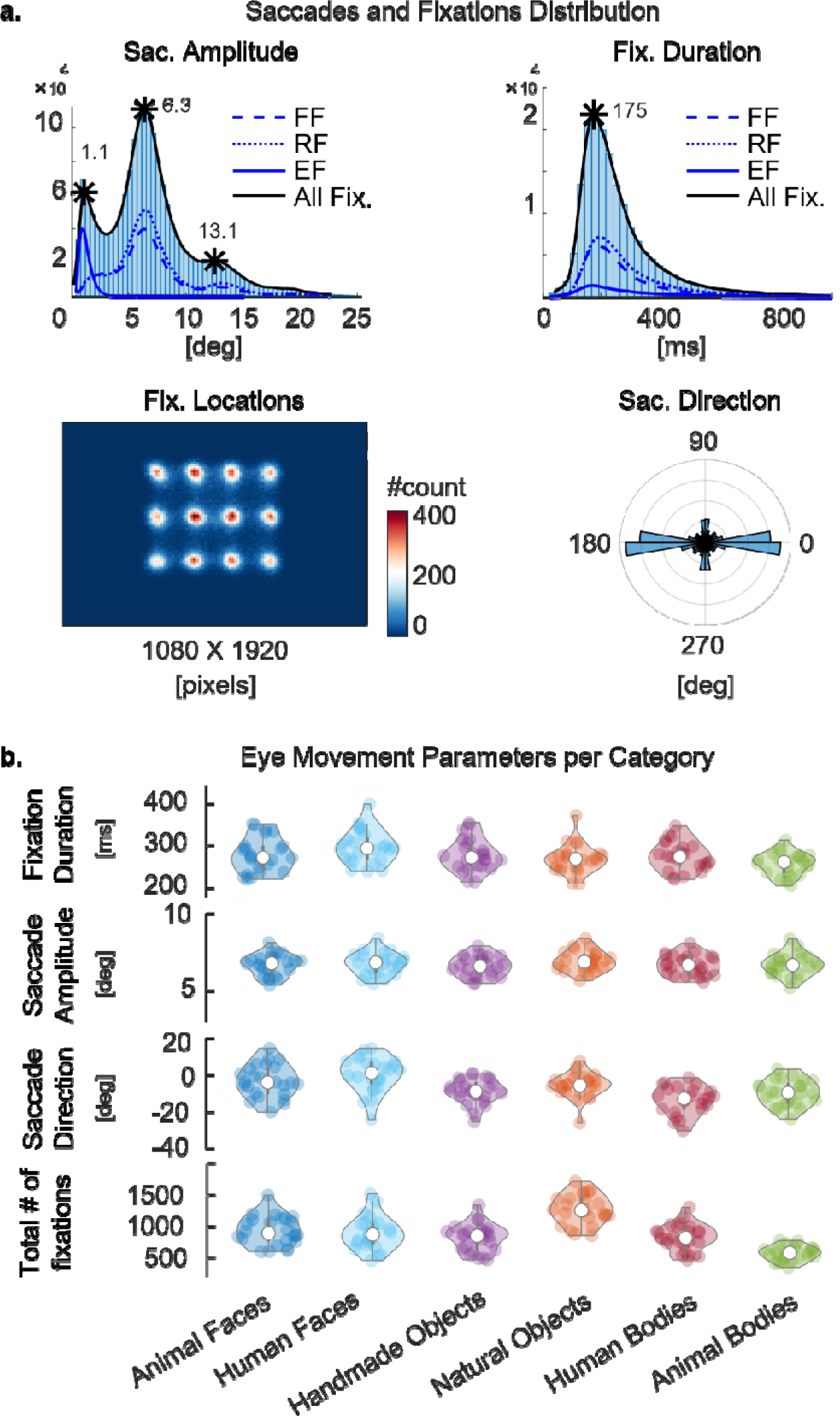
Behavioral results and eye movement properties. **a.** Saccade and fixation distributions for data combined over all subjects (n=174,648 fixations). Top-left: saccade-amplitude distribution with three peaks reflecting the search array structure. Bottom-left: heatmap of the spatial distribution of number of fixations across the screen. Top-right: ex-Gaussian distribution of fixation durations. Bottom-right: angular histogram of saccade directions showing a clear horizontal bias in the search direction. **b.** Eye movement parameters for each category separately. Saccade-direction is relative to a right horizontal direction (see panel a). For violin plot notations, see Figure 1.

#### Univariate Analyses

To examine the univariate (individual electrodes) regression ERP/FRPs, single trials from the training set of one of the folds were low-pass filtered with a cutoff of 30 Hz using a 4th-order non-causal Butterworth filter (we used only the training set, i.e 75% of the data, so that we present the same data used later for training the MVPA model; similar results are obtained with the full dataset). For each subject separately, we averaged the single trials (−100 to 700 ms post-fixation \ stimulus onset). We examined category-related differential activity in three contrasts: Human Faces - Inanimate Objects (combining HO and NO categories), Human Faces - Animal Faces, and Animate (combining HF, AF, AB, HB) - Inanimate Objects (Figure 1.c). Prior studies, including our previous FRP study, typically found maximal categorical selectivity, especially for faces, in right parieto-occipital electrodes, like the P10, around the latency of the visual N1 (Bentin et al., 1996; Itier and Taylor, 2004; Rossion, 2014; Auerbach-Asch et al., 2020). We therefore apriori selected electrode P10 for the univariate analysis.

We tested for significant differences between categories using a cluster permutation test (Maris and Oostenveld, 2007). For this test, a two-tailed t-test was conducted for each time point, and clusters were defined by temporally adjacent points that crossed a threshold of p<0.01. Points were merged into a cluster only if they were both negative or both positive. The cluster statistic was defined as the absolute sum of t-values (t_sum) of all points in the cluster. We constructed a null distribution of cluster statistics by repeating the procedure after randomly permuting the assignment of category labels (1000 permutations). From each iteration, we extracted the largest t_sum and included it in the null distribution. Finally, clusters in the real data with t_sum>95 percentile of the null distribution were considered significant (Figure 4).

**Figure 4.**
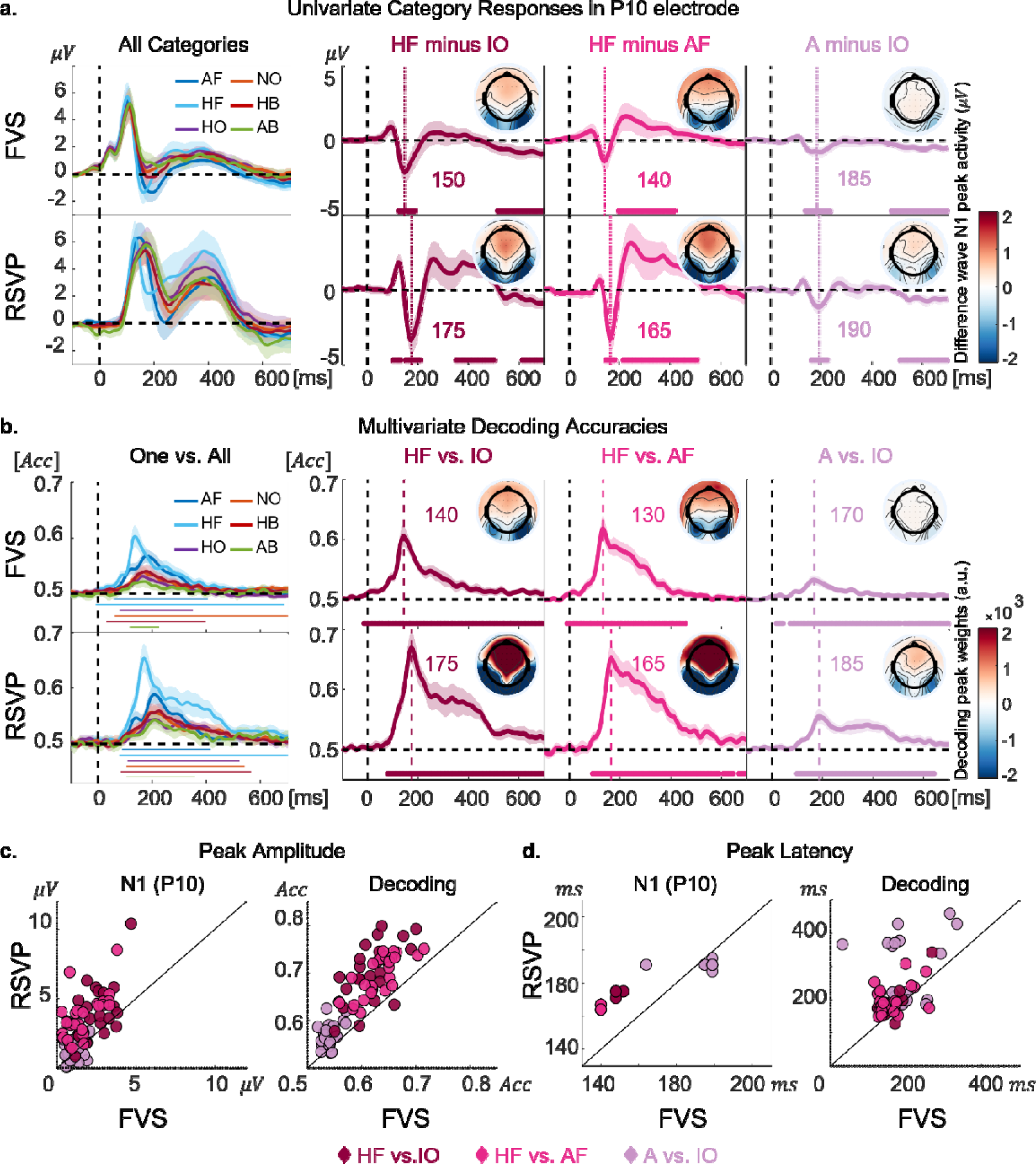
Univariate and multivariate category information. **a-b.** Univariate (**a**) and multivariate (**b**) results analysis. Top rows: FVS data, time-locked to self-driven fixations. Bottom rows: RSVP data, time-locked to stimuli onsets. Left Column: grand-average results (across subjects) in electrode P10 (**a**) and a one-vs-all decoding design (**b**), for all six categories. Right three columns: Grand-average results (across subjects) for three categorical comparisons of interest. Vertical dotted lines mark N1 peak latencies in electrode P10 in **a**, and peak decoding latencies in **b**. The insets show scalp topographies in a 20 ms window around the peak latency which is specified next to each inset. The shaded lines represent 95% confidence intervals across subjects. The horizontal bars below each graph mark time points in which the differential activity (**a**) or decoding accuracies (**b**) were significantly different from zero (**a**) or 0.5 chance level (**b**) by cluster permutations (p < 0.05). **c-d.** Comparison of peak amplitude (**c**) and peak latency (**d**) between the two viewing conditions. Left: univariate differential category response absolute peak value. Right: multivariate decoding peak accuracy. Empty circles depict single-subject data and filled circles depict the mean across subjects (both color-coded by category comparison). AF - Animal Faces; HF - Human Faces; HO - Handmade Objects; NO - Natural Objects; HB - Human Bodies; AB - Animal Bodies, A - Animate, IO - Inanimate Objects

We detected single-subject peak category-related activity (maximal amplitude of the contrast difference wave) during the N1 component latency window (between 110 to 210 ms), separately for each viewing condition. To determine if these were influenced by viewing condition, we conducted two-tailed, paired t-tests for the three categorical comparisons (HF-IO; HF-AF; A-IO) and corrected our p-value threshold to p<0.017 (Bonferroni correction accounting for the three comparisons). Finally, Cohen’s effect-size value was calculated by 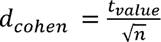 for paired t-tests (Rosenthal and Rosnow, 2008).

To analyze the differential N1 peak latency, we used a jackknife procedure, which was shown to be robust for latency analysis (Miller et al., 1998). Following this procedure, 24 sub-averages of activity in electrode P10 were calculated, each time leaving one subject out of the average. We detected the peak latency of the sub-averages in the N1 time range and again conducted separate two-tailed t-tests to test for differences between viewing conditions for each categorical comparison. We statistically corrected the t values obtained using the jackknife procedure by dividing the t values by the number of subjects (Kiesel et al., 2008), and used a critical p<0.017 accounting for the 3 latency measures. Cohen’s-d effect size was calculated on the jackknife-corrected t-values.

#### Multivariate-Decoding Analyses

Having obtained a time-point by time-point decoding accuracy measure for each subject, we tested for significant decoding accuracy using a one-tailed, cluster-based permutation test across subjects (1000 permutations) against a 0.5 chance baseline. As with the univariate analysis, points inclusion in clusters were selected by a conservative threshold of p<0.01. The t-distribution under the null hypothesis was built by randomly permuting labels of the actual accuracies and chance baseline. Finally, clusters in the real data with a t_sum value larger than 95% of the null distribution were defined as significantly different from chance.

We compared peak decoding accuracies between the RSVP and FVS conditions using two-tailed, paired t-tests for the three categorical comparisons (HF vs. IO; HF vs.AF; A vs.IO) and corrected our p-value threshold to p<0.017 (Bonferroni correction accounting for the three comparisons). To compare the temporal dynamics of categorical information in FVS and RSVP, we tested for significant differences between the viewing conditions in the decoding onset and peak latencies. As for the univariate analysis, we used a jackknife procedure (Miller et al., 1998; Ulrich and Miller, 2001; Kiesel et al., 2008), by computing 24 grand-average waveforms from a subsample of n-1 participants such that each participant was omitted once from the subsample average. On each subsample average, we detected the latency of the peak accuracy. We computed the decoding onset on each subsample in two ways (for robustness): (1) The first time point for which False Discovery Rate (FDR)-corrected p-values were smaller than 0.05 (Benjamini and Yekutieli, 2001), for at least 50 consecutive milliseconds, in a time-point by time-point one-tailed t-tests against chance (we refer to this as the Above Chance FDR onset latency); (2) The first time point for which decoding accuracy crossed a threshold defined as the mean plus two standard deviations of the baseline decoding accuracy (−200 to −50 ms), provided that the consecutive 50 ms met the same criterion (we refer to this as the Baseline Threshold onset latency). This procedure resulted in three latencies for each viewing condition: peak latency and two onset latencies.

To test the hypothesis that decoding onset latencies are different between viewing conditions, we conducted a two-tailed t-test with jackknife-corrected t-values and Bonferroni correction to account for the three category comparisons.

Finally, an analysis of the differences in the across-condition and within-condition peak decoding latencies was conducted by detecting the two-dimensional peak (latency_train, latency_test) in the temporal generalization matrices (see MVPA Decoding). The two-dimensional TGMs were smoothed using a 10-sample square kernel prior to identifying the peak to avoid spurious peaks. We then calculated the difference *D_l_*between the latency of the peak on the train and test axes:

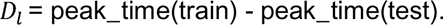

Within a condition, we expect *D_l_* to be zero (i.e. the 2-dimensional peak would fall on the diagonal of the TGM). Across conditions, negative *D_l_* would indicate that the decodable information evolves earlier in the condition used for training compared to the condition used for the testing and vice versa if *D_l_* is positive. A two-way repeated-measures ANOVA across subjects was applied to the peak-latency difference values (*D_l_*), with the factors Decoding Method (within, across) and Viewing Condition (FVS, RSVP).

## Results

### Behavior

Twenty-four participants performed a dual task, including target detection (flowers or butterflies) and recognition memory (Figure 1) in sets of 12 objects from six categories (Figure 1b-c), which were shown in one of two viewing conditions (Figure 1a). In the FVS condition, all 12 objects were presented simultaneously in an array and participants actively searched the object arrays at a self-driven pace. In contrast, in the RSVP condition, participants passively viewed the objects presented serially at the center of the screen. Performance in both target detection (accuracy mean±SEM: FVS: 0.92±0.009, RSVP: 0.92±0.007) and recognition memory (FVS: 0.8±0.01, RSVP: 0.75±0.01) was well above chance for both viewing conditions (Figure 1d). Recognition memory performance was higher in the FVS condition. A two-way ANOVA confirmed a main effect of Task (target detection better than recognition *F*(1,23) = 207.4; *p* = 5.35 10^−13^, *generalized eta square (ges)* = 0.61), a main effect of Viewing Condition (FVS better than RSVP, *F*(1,23) = 6.9, *p* = 0.01, *ges* = 0.07), and a significant interaction (*F*(1,23) = 21.4, *p* = 1.2 10^−4^, *ges* = 0.1). Follow-up contrasts showed that the difference in performance between viewing conditions was significant in the recognition task (*t*(47) = 5, *p* = 6.5 10^−6^, *d_Cohen_* = 1.02) but not in target detection (*t*(47) = −0.6; *p* = 0.5; Figure 1d). The better recognition memory in the FVS compared to RSVP condition likely stems from the fact that the subjects could visit objects more than once and dwell for longer periods (Zelinsky et al., 2011; Meghanathan et al., 2019), as well as actively control the information acquisition (Markant et al., 2016).

### Eye Movements

In the FVS condition, objects were displayed in a 3X4 array, such that each object was of size 3°X3° and object centers were separated by 6° (Figure 1a). A total of 7,486±340 (mean±SEM across subjects) fixations per subject were detected in the FVS condition. The distribution of saccade sizes, showing three prominent peaks (∼1.1°, ∼6.3°, and ∼13°), together with the heatmap of all fixation locations during FVS, indicate that participants either made saccades between objects (FF\RF) or within objects (EF), and rarely dwelled in spaces between objects (Figure 3a). The angular histogram of saccade directions indicates that participants mainly scanned the stimuli along the horizontal axis (as in Gilchrist and Harvey, 2006). Overall fixation durations exhibited a typical ex-Gaussian distribution with a peak at 175 ms (Figure 3a). On average, in each FVS array, subjects fixated on 10.8±0.14 (mean±SEM) objects (out of 12) at least once, and made 13.5±0.9 (mean±SEM) return fixations (RF). Figure 3b displays eye-movement parameters for each of the six object categories separately (Figure 1c). These parameters were accounted for in obtaining GLM-reconstructed single trials (see <u>Temporal Signal Deconvolution using GLM</u>).

### Similar Category-Specific Information in Passive and Active Vision

#### Univariate Analysis

Before delving into multivariate analyses, which were the main goal of the study, we verified that univariate (individual electrodes) rFRPs to images of different categories, time-locked to fixation onsets, resemble those expected from previous rERP and rFRP studies. Figure 4a depicts the responses to the different categories in the two viewing conditions at the P10 electrode, where previous ERP studies showed maximal category-selective responses (Itier and Taylor, 2004; Rossion and Jacques, 2011). We focus on three planned categorical comparisons driven by the literature: Human Faces vs. Inanimate Objects (HF vs. IO; Bentin et al., 1996), Human Faces vs. Animal Faces (HF vs. AF; Kaneshiro et al., 2015), and Animate vs. Inanimate Objects (A vs. IO; Kriegeskorte et al., 2008; Cichy et al., 2014). All three differential responses show posterior negativity and frontal positivity during the differential N1, in both active (FVS) and passive (RSVP) conditions. The peaks of absolute differential responses were larger in ERPs (RSVP) relative to FRPs (FVS) for the HF-IO comparison (*t*(23) = 4.9, *p* = 5.7 10^−5^, *d_Cohen_* = 1), In line with our previous findings (Auerbach-Asch et al., 2020), and similarly for the HF-AF comparison (*t*(23) = 8.3, *p* = 2.3 10^−8^, *d_Cohen_* = 1.7; Figure 4c). The A-IO comparison showed the same trend, but it did not survive the Bonferroni multiple-comparisons correction (*t*(23) = 2.2, *p* = 0.04, *d_Cohen_* = 0.45; Figure 4c). Unlike our previous study, peak latencies were earlier in FVS condition compared to RSVP condition for HF-IO comparison (*t*(23) = 3.7, *p* = 0.001, *d_Cohen_* = 0.75) and HF-AF (*t*(23) = 9.1, *p* = 9.1 10^−4^, *d_Cohen_* = 0.77), but not for A-IO comparison (*t*(23) = −0.2, *p* = 0.8, *d_Cohen_* = −0.04; Figure 4d). Since the distribution of fixation durations in the FVS condition includes infrequent long fixations, not matched by SOAs in the RSVP condition, we repeated the FVS analysis using only fixation durations between 100-300 ms (matching SOA distribution in the FVS), which yielded qualitatively similar results.

#### Multivariate Analysis

MVPA enables us to extend our analysis from predefined electrodes of interest and the mean activity over events, to information (operationalized as above-chance decoding) available across all electrodes at the single event (fixation or stimulus onset) level. All six categories could be significantly decoded from EEG activity locked to single fixations during unrestricted active viewing. (Figure 4b). Human faces yielded the highest decoding accuracies. The topography of the classifier weights (corrected to the forward model according to; Haufe et al., 2014); Figure 4b insets) at the peak decoding latency resembled the topographies of the peak FRP\ERP differential activity (Figure 4a-b insets). Similar to the univariate results, peak decoding accuracies were significantly higher in the RSVP compared to the FVS condition for the three categorical comparisons (*HF vs. IO*: *t*(23) = 8.2, *p* = 2.8 10^−8^, *d_Cohen_* = 1.7; *HF vs. AF*: *t*(23) = 10.5, *p* = 2.8 10^−10^, *d_Cohen_* = 2.1; *A vs. IO*: *t*(23) = 6, *p* = 3.7 10^−6^, *d_Cohen_* = 1.2; Figure 4c). Decoding peak latencies were shorter during FVS compared to RSVP (Figure 4d), a finding that is elaborated on in the following section.

### Categorical Information Emerges Earlier in Active Vision Compared to Passive Vision

Having shown that categorical information is decodable in both constrained and free viewing, we next compared the dynamics of categorical information. We first asked if information evolves with the same dynamics across viewing conditions, addressing both the onset and peak latency of decoding. For robustness, we determined the decoding onset latency in two complementary ways: Baseline Threshold, and Above Chance FDR (see <u>Multivariate-Decoding Analysis</u> section for details). Overall, category information in the FVS condition emerged earlier than in the RSVP condition (Figure 5). Peak decoding latencies were significantly earlier in the FVS compared to the RSVP condition for HF\O, and HF\AF contrasts (Table 1, left column). Onset latencies were significantly earlier in the FVS condition than in the RSVP condition in all comparisons for at least one of the onset estimates (Table 1, middle and right columns; note that the Bonferroni correction is highly conservative here, considering that the tests are not completely independent).

**Figure 5.**
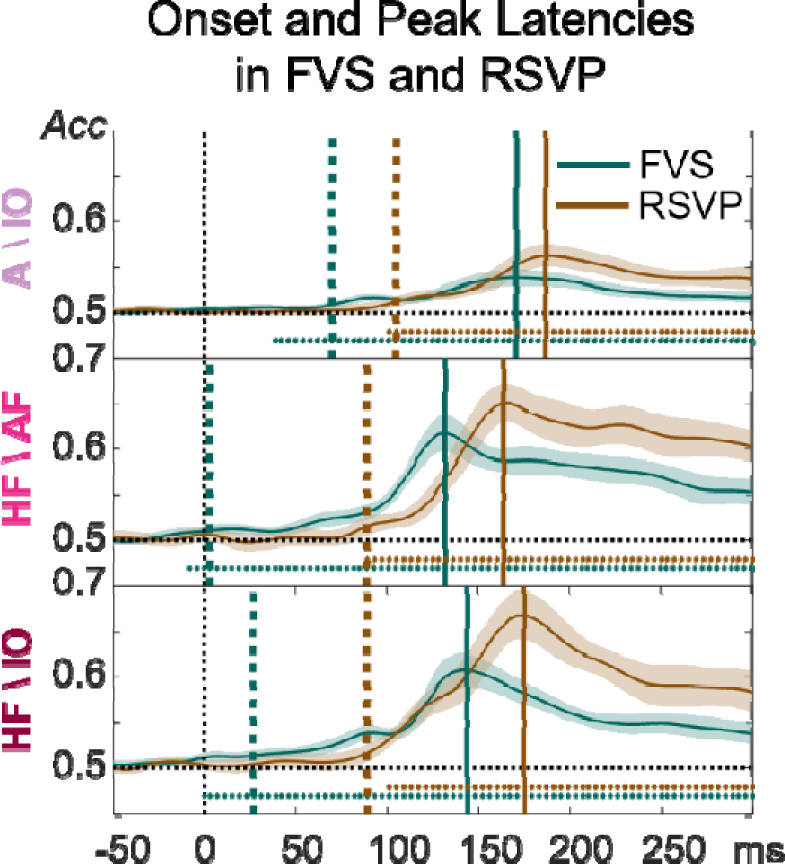
Decoding dynamics of FVS compared to RSVP. Active (FVS) and passive (RSVP) decoding accuracies for the three categorical comparisons and three latency measures. Solid vertical colored lines depict the latency of the grand-average decoding peak. Dotted vertical colored lines depict the decoding onset latency determined using the Baseline Threshold criteria. The horizontal dotted lines depict the time points in which decoding accuracies were significantly above chance, after correcting for multiple comparisons using False Discovery Rate (FDR); The first significant time point of the cluster is the Above-Chance FDR Onset; see Methods section <u>Multivariate-Decoding Analyses</u>). All three measures show earlier latencies for the FVS condition.

**Table 1.**
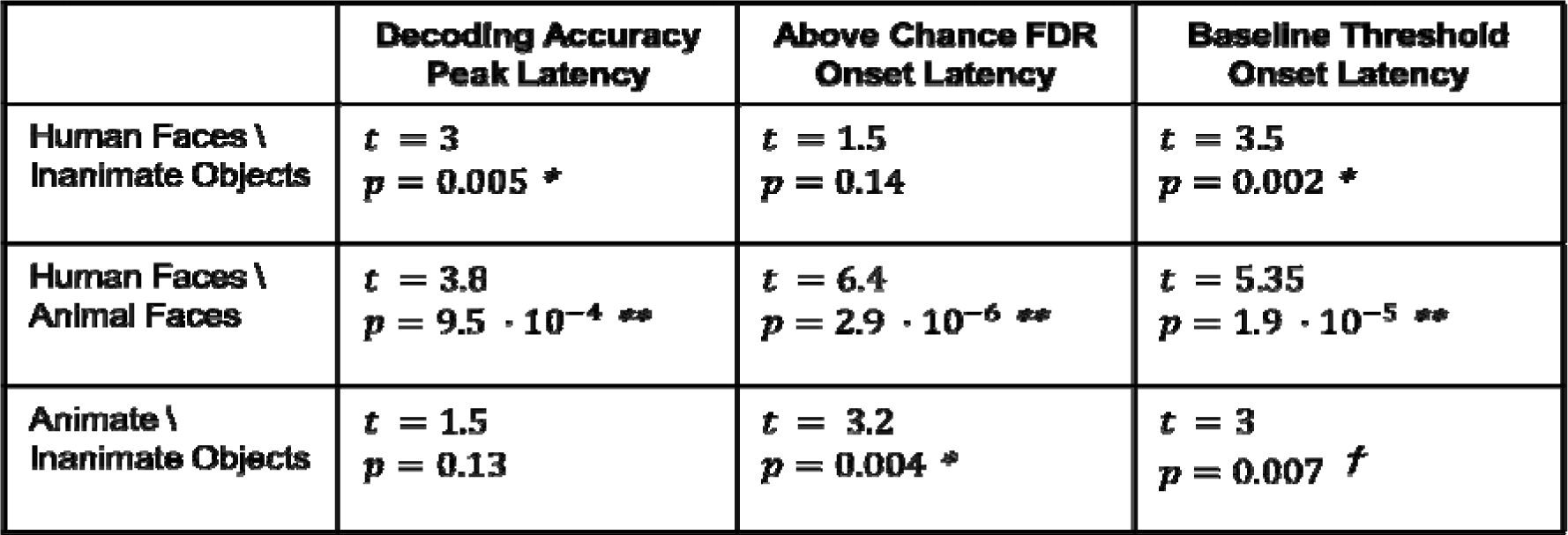
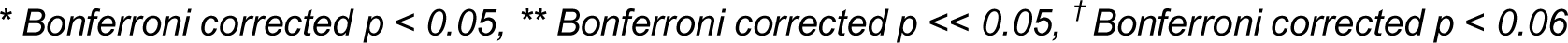
Statistical comparison of FVS and RSVP onset and peak latencies. Comparison between the active (FVS) and passive (RSVP) viewing conditions considering peak decoding latency and two measures of decoding onset. T-values and p-values are corrected for the jackknife procedure. Symbols denote significance following additional Bonferroni correction for nine comparisons.

|  | **Decoding Accuracy**<br>**Peak Latency** | **Above Chance FDR**<br>**Onset Latency** | **Baseline Threshold**<br>**Onset Latency** |
| --- | --- | --- | --- |
| **Human Faces \**<br>**Inanimate Objects** | $t = 3$<br>$p = 0.005$ * | $t = 1.5$<br>$p = 0.14$ | $t = 3.5$<br>$p = 0.002$ * |
| **Human Faces \**<br>**Animal Faces** | $t = 3.8$<br>$p = 9.5 \cdot 10^{-4}$ ** | $t = 6.4$<br>$p = 2.9 \cdot 10^{-6}$ ** | $t = 5.35$<br>$p = 1.9 \cdot 10^{-5}$ ** |
| **Animate \**<br>**Inanimate Objects** | $t = 1.5$<br>$p = 0.13$ | $t = 3.2$<br>$p = 0.004$ * | $t = 3$<br>$p = 0.007$ † |

\* Bonferroni corrected $p < 0.05$ , \*\* Bonferroni corrected $p < 0.01$ , <sup>†</sup> Bonferroni corrected $p < 0.06$

Earlier decoding onsets in the active FVS condition may indicate one of two alternatives: either information is represented in a similar way (similar topographical distinctions) across viewing conditions but evolves earlier during the active condition, or the categorical representation is not only earlier, but also different between viewing conditions. To test this, we performed cross-decoding of categorical information when training the classifiers on data from the passive condition and testing them on the active condition (and vice versa), applying temporal generalization analysis (King and Dehaene, 2014). Figure 6a depicts the mean TGMs across subjects for the categorical classification of HF vs. IO, HF vs. AF, and A vs. IO. Each data point in a TGM depicts the decoding accuracy based on a classifier trained for a given time point and tested on separate data at the same (diagonal) or another (off-diagonal) time point. In across-condition TGMs, the classifiers were trained on data from one viewing condition and tested on data from the other. In within-condition TGMs, the training and testing were performed using a cross-validation procedure. The results show significant above-chance decoding in the across-condition, suggesting shared representations in the two viewing conditions (Figure 6a).

**Figure 6.**
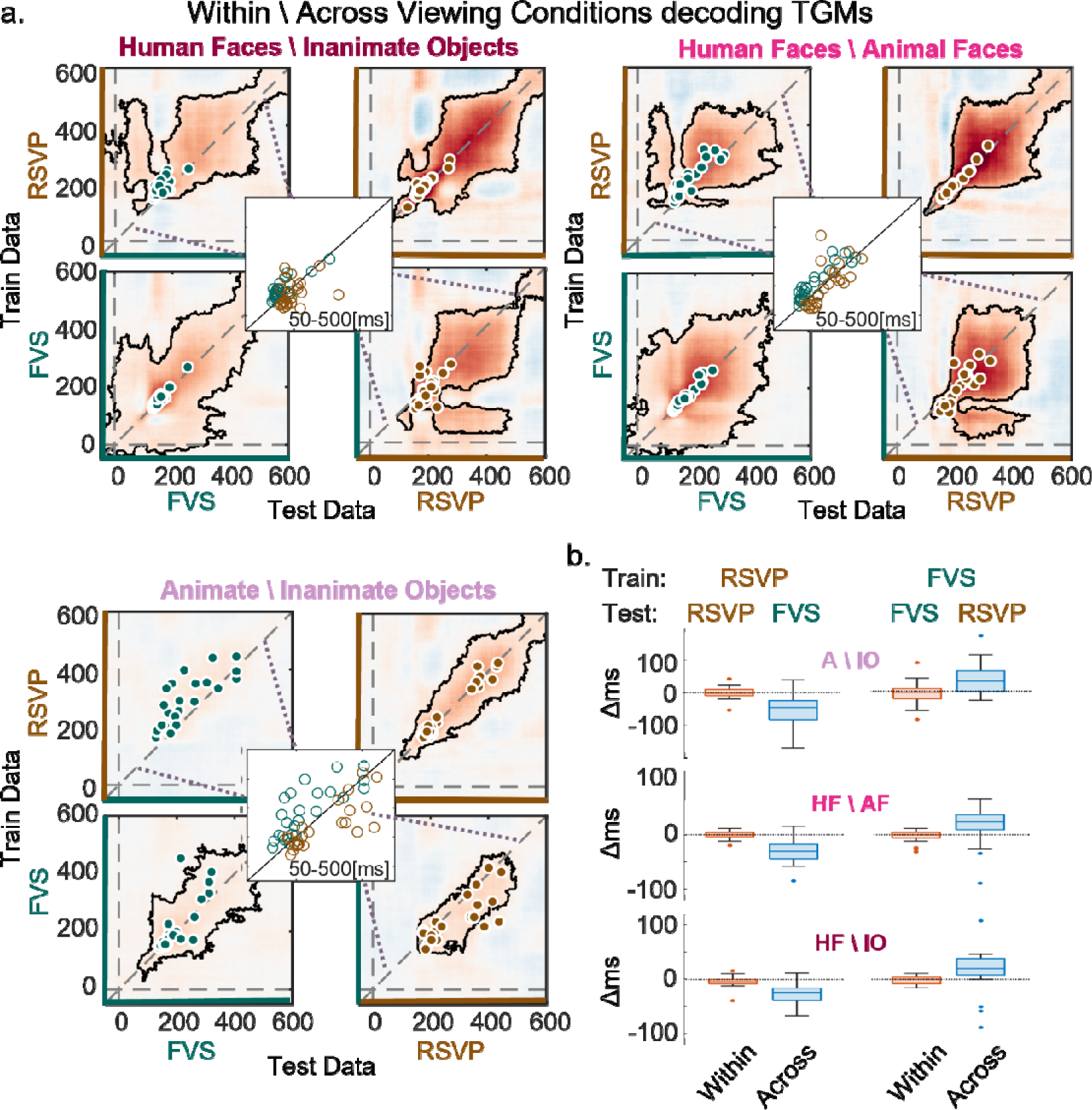
Within and Across-condition decoding. **a**. Temporal Generalization Matrices showing decoding accuracies of classifiers trained separately on each time point (y-axis) and tested on all time points of both viewing conditions (x-axis). For each categorical comparison: rows depict the train-set condition (top: RSVP, bottom: FVS), and columns depict the test-set condition (left: FVS, right: RSVP). Dots represent single-participant latencies of the two-dimensional peak in decoding accuracy (green for FVS test sets and brown for RSVP test sets). The black outline depicts the two-dimensional clusters of significant decoding accuracies (across participants). The center inserts overlay the two cross-decoding peak accuracies to show the shift in latencies - brown: train on FVS, test on RSVP, green: train on RSVP, test on FVS. **b.** Distance of the two-dimensional decoding peak latency from the diagonal, reflecting the difference in training and testing latencies of the peak decoding accuracies of the TGMs in a. The box represents the interquartile range (IQR) between the 0.25 and 0.75 quantiles. The horizontal line within the box is the sample median. The dots outside the box are the outliers which are more than 1.5*IQR away from the box edges. The vertical lines exceeding the box limits connect the maximum and minimum non-outlier values in the data.

The level of symmetry across the diagonal of across-condition decoding TGMs indicates whether the dynamics of the representation are similar or shifted in time. If the representational dynamics are similar in format and temporal progression, decoding accuracies would be symmetric over the diagonal, and the peak decoding would fall along the TGM diagonal. In contrast, if representations are shared in format, but shifted in time, the peaks of cross-condition decoding accuracy will lay off-diagonal. The results clearly supported the latter case. Individual subjects’ decoding peaks (marked by colored dots on the TGMs in Figure 6a) lay on the diagonal in the within-condition classifiers, as expected. Conversely the cross-condition decoding peaks lay above the diagonal (t_train > t_test) when training on RSVP and testing on FVS, and below the diagonal (t_train < t_test) when training on FVS and testing RSVP, indicating earlier times during the FVS compared to RSVP condition; see center insets of Figure 6a). To confirm this observation statistically, we ran a two-way ANOVA with factors Train-set Condition (FVS, RSVP) and Decoding Method (within, across viewing conditions) with the dependent variable being the difference between training and testing latencies at peak decoding (proportional to the distance from diagonal). The ANOVA confirmed a main effect of Train-set Condition (*F*(1,23) = 16.3, *p* = 5.2 10^−4^, *ges* = 0.21), qualified by a significant interaction between the Train-set and Decoding-Method factors (*F*(1,23) = 15.4, *p* = 6.710^−4^, *ges* = 0.2). Figure 6b shows that this interaction is due to a delayed peak decoding latency when training on the FVS and testing on RSVP and an earlier peak decoding latency when training on RSVP and testing on FVS. Taken together, we conclude that category representations are shared between active and passive viewing, yet evolve earlier in active vision.

## Discussion

Visual processing in the brain is typically measured time-locked to abrupt onsets of external stimuli, while constraining self-driven eye movements. Therefore, the evidence primarily focuses on reactive processes rather than proactive ones based on perceptual and temporal predictions (Bar, 2004; Kersten et al., 2004; Clark, 2013). Here, we tracked category representation dynamics with EEG activity time-locked to eye fixations in self-paced visual search, comparing it to representation dynamics time-locked to stimuli onsets, in rapid serial visual presentation. Our results show that despite the methodological complexity of FVS paradigms, category information can be decoded from single fixations on objects using multivariate pattern analysis. Cross-condition decoding showed that object-category representations are largely shared between active and passive viewing conditions, supporting the generalizability of models based on traditional gaze-controlled paradigms. However, we show that categorical representations evolve (and therefore can be decoded) faster when participants actively sample visual information rather than obtain it passively.

Previous studies using MVPA on EEG data show that object category representations can be decoded from single trials during passive-viewing conditions (Simanova et al., 2010; Carlson et al., 2011, 2013; van de Nieuwenhuijzen et al., 2013; Cichy et al., 2014; Haxby et al., 2014; Kaneshiro et al., 2015; Contini et al., 2017; Dobs et al., 2019; Grootswagers et al., 2019b; Robinson et al., 2019). Representations of human and animal faces were dissociable in these studies, both being highly discriminable from other categories. Whereas our results replicate these findings in the passive condition and generalize them to the more natural active viewing, the features upon which two categories are successfully discriminated could be different in the two viewing conditions. However, across-condition decoding resulted in significant accuracies indicating largely shared representations when viewing objects actively or passively. Taken together, the results simultaneously support the ecological validity of controlled serial visual presentation, and the feasibility of the MVPA analysis in the context of noisy free-viewing experiments, opening the way to studies involving more ecological free viewing situations.

While activity patterns were similar across conditions, univariate and multivariate analyses both revealed that information evolves ∼25-30 ms earlier, and is attenuated, in active compared to passive vision. Accordingly, in the cross-decoding analysis, time-point-specific classifiers trained on the active condition data best decoded data from later times in the passive condition, and vice versa. The results thus indicate that shared object representations emerge earlier when obtained actively.

What could be the causes for the attenuated and earlier representations in active-compared with passive viewing? A core difference between the conditions is the augmented ability to form sensory, semantic, and temporal expectations during active viewing. First, memories from previously visited locations in the scene may constrain perceptual hypotheses about the next observed stimulus. Second, parafoveal information, although impoverished, may provide information on the content of the next possible fixation targets (Dimigen et al., 2012; Kornrumpf et al., 2016; Niefind and Dimigen, 2016; de Lissa et al., 2019; Huber-Huber et al., 2019; Buonocore et al., 2020). Third, internal action-related signals inform sensory systems of the upcoming trajectory, timing, and sensory consequences of eye movements (Brown et al., 2013; Cavanaugh et al., 2016; Sun and Goldberg, 2016; Subramanian et al., 2019).

There is a wealth of evidence for the attenuation of neural responses when information is expected (interpreted as reduced prediction error; for a review see (de Lange et al., 2018). This could explain the larger univariate effects in the passive condition, in which neither temporal nor categorical stimulus properties can be expected, compared to the active condition. As for the effect of expectations on decoding, previous results are mixed. Some studies have shown that motor or perceptual expectations lead to higher stimulus decoding accuracies (Brandman and Peelen, 2017; Yon et al., 2018). Such results support the sharpening of representations as an account for expectation suppression, by which highly selective neurons are activated by expected stimuli while weakly selective neurons are silenced (Kok et al., 2012). In contrast, other studies have found either similar decoding accuracies for expected and unexpected stimuli or even higher decoding accuracies for unexpected, rather than expected, stimuli (Kumar et al., 2017; Richter et al., 2018), supporting the dampening account of expectation-suppression by which highly selective neurons respond less to the expected stimuli, reducing signal-to-noise ratios (Kersten et al., 2004; Murray et al., 2004). Our results are consistent with the latter studies. However, the free-viewing condition is intrinsically more variable than the highly controlled RSVP, including parallel processing of parafoveal objects, variable gaze positions, and oculomotor-induced noise in the EEG measurement. Thus, we cannot rule out the possibility that the lower decoding accuracies in FVS are a result of such methodological issues.

In contrast to expectation effects on response amplitudes, there is less evidence, and less theoretical discussion, regarding the expectation effects on the temporal evolution of neural representations, or the dynamics of prediction error processing. The current results indicate earlier evolution of categorical information in active viewing, during which temporal and categorical expectations can be formed. According to the predictive coding framework, recognition is achieved as predictions are updated in the process of minimizing prediction errors (Friston, 2010; Hohwy, 2012; Clark, 2013). Categorical expectations are based on contextual cues obtained during the scanning (Yan et al., 2023), parafoveal information (Buonocore et al., 2020; Huber-Huber et al., 2021), and even gradual release from saccadic suppression before the end of the saccade (Diamond et al., 2000; Bremmer et al., 2009). All these factors may lead to faster convergence of the recognition process. Indeed, in our previous study which, by design, precluded parafoveal viewing, fixation-related face-effects peaked slightly later, not earlier, than the face-effect in an ERP condition (Auerbach-Asch et al., 2020). It is also possible that processes related to the motor plan (corollary discharge) promote anticipatory categorical processing, by pre-saccadic remapping of sensory information, during which neurons respond to information present in the saccade target location, outside their classic receptive field (Duhamel et al., 1992; Melcher, 2007; Crapse and Sommer, 2008; Rao et al., 2016). In addition, pre-saccadic spatial attention shifts may result in higher sensitivity of neurons whose receptive field corresponds to the saccade target location (Carrasco, 2011; Rolfs and Carrasco, 2012). Nevertheless, we did not observe pre-saccadic category information, nor did Fabius et al., 2020, in the context of decoding spatial frequencies. Such pre-saccadic information might be easier to detect using more sensitive measures than M\EEG (e.g. Electrocorticography; Dürschmid et al., 2016; Boring et al., 2020).

The previous discussion focused on expectations regarding the category of the stimulus. An additional explanation for hastened dynamics in active vision could be expectations about the precise timing of incoming information (Nobre et al., 2007). Temporal expectation is thought to ensure optimal neural preparation at the right moment (Rohenkohl and Nobre, 2011). This leads to enhanced behavioral speed and accuracy (Correa et al., 2005; Nobre et al., 2007; Rolke and Hofmann, 2007), and higher amplitude of neural response to stimuli at expected times (Nobre and van Ede, 2018). Less is known about the effect of temporal expectations on the temporal dynamics of post-stimulus neural processing. Van Ede et al., 2018, found increased accuracy in decoding the orientation of temporally anticipated gratings, but no effect of expectation on the decoding onset latency. However, temporal expectations in that study were based on external cues unrelated to self-generated action signals. A recent study, van den Brink et al., 2021, found that temporal predictions significantly hastened the onset latency of the Centroparietal Positivity (CPP) ERP component, considered to be a neural signature of decision formation. Whether the hastening of decision onset is due to speeded sensory processes or strategic top-down effects was left unanswered. Our results indicate the faster formation of perceptual representations during active vision, much earlier than the process indicated by the CPP.

Taken together, active and passive viewing conditions share similar perceptual representations with different temporal dynamics. Thus, the representational space found in passive viewing studies holds also for more natural conditions. The accelerated dynamics of information processing in active vision elucidates the role of active sensing in shaping perception.

### Methodological Limitations and Considerations

The definition of saccades and fixations onset latencies may depend on the specific algorithm used for event detection from eye tracking data (Andersson et al., 2017; Birawo and Kasprowski, 2022). If the widely used Engbert and Mergenthaler algorithm (EMA; Engbert and Mergenthaler, 2006) that we employed introduced a systematic delay in estimated fixation onset latencies, it could explain the apparently earlier decoding times in the FVS condition. However, the EMA tends to produce relatively shorter saccade durations, entailing earlier, rather than later, fixation onsets, compared to other algorithms. Additionally, the 8 ms difference between the algorithm producing the shortest saccade durations and the EMA in a recent review (Andersson et al., 2017), cannot explain the ∼30 ms difference in category representations we found between active and passive viewing conditions. Finally, in our previous study using FRPs, categorical information emerged later in FRPs than ERPs when using either the Eyelink proprietary algorithm or the EMA, supporting the conjecture that the stimuli and paradigm determine the latency differences, not the choice of algorithm (Auerbach-Asch et al., 2020).

Deconvolution methods are used to distinguish co-occurring influences of information processing and overlapping signals (Ouyang et al., 2015; Ehinger and Dimigen, 2019). In theory, given enough variability in the data, machine learning methods such as MVPA time-resolved decoding are also capable of distinguishing such responses without the need for prior temporal deconvolution (Grootswagers et al., 2019a, 2019b, 2022; Robinson et al., 2019). However, this depends on balanced designs and the independence of sequential events, which are hard to control in free viewing (e.g., category ‘A’ being viewed more frequently after ‘B’ compared to other categories). Thus, the initial deconvolution of overlapping activity, where information about the timing of fixations and the content viewed is used, may better control for sequential effects (Petruo et al., 2021).

The stimulus set used in this study has been extensively studied by others (Kiani et al., 2007; Carlson et al., 2013; Cichy et al., 2014; Kaneshiro et al., 2015; Grootswagers et al., 2017). Repeated use of this set may introduce some bias or lack of generalizability (Grootswagers and Robinson, 2021), yet has the advantage of comparability between imaging methods (Kriegeskorte et al., 2008; Cichy et al., 2014), and establishing replicability. In particular, it was advantageous in our study, as this is the first attempt to establish the decodability of object category representations during active vision.

Our study used arrays of isolated items without context. Context provides even more constraints to, and better prediction of, the possible identity of objects in a scene (Mudrik et al., 2010; Brandman and Peelen, 2017). If the acceleration of representation we observed is based on the semantic predictability of objects, we predict even earlier decoding times when objects are presented in context. However, if this acceleration is solely due to temporal predictions, then the same temporal dynamics are expected in random arrays and coherent scenes.

## Conclusions

Object recognition during natural vision is accomplished by actively sampling the visual information in a series of fixations. Here, we showed that object categories can be decoded from EEG activity locked to single fixations. Cross-condition decoding revealed that categorical representations are largely shared between active and passive viewing, supporting the generalizability of models based on traditional gaze-controlled paradigms. However, categorical representations evolve more quickly when participants actively sample visual information rather than obtain it passively. We attribute these faster dynamics to content and temporal expectations available when subjects can actively control the input, rather than passively observe externally evoked stimuli.

## Acknowledgments

This work is supported by grant 1902/14 from the Israel Science Foundation to L.Y.D. We thank Dr. Edden Gerber for his MATLAB implementation of the preprocessing procedure. We thank Chen Gueta and Gal Chen for their help in statistical analysis, all the research assistants for their help in data collection, and all the Deouell-lab members for their valuable comments.

## Footnotes

1 Since the focus of this study is on representations of object categories, to prevent confusion we replaced the commonly used notation *cat* for categorical predictors in GLM models with *bin,* indicating that the value of these predictors was either 0 or 1.

